# *Pantoea ananatis*-triggered systemic resistance requires root sensing through the LORE receptor kinase in Arabidopsis

**DOI:** 10.1101/2025.11.18.688997

**Authors:** Simon Duchateau, Jérôme Crouzet, Sylvain Cordelier, Matthieu Touchard, Romain Schellenberger, Célia Borrego, Sandra Villaume, Jean-François Guise, Qassim Esmaeel, Marie-Christine Groleau, Maude Cloutier, Charles Gauthier, Sandrine Dhondt-Cordelier, Florence Mazeyrat-Gourbeyre, Fabienne Baillieul, Stefanie Ranf, Eric Déziel, Aziz Aziz, Stéphan Dorey

**Affiliations:** University of Reims Champagne-Ardenne, RIBP-USC INRAE 1488, 51100 Reims, France; INRS-Centre Armand-Frappier Santé Biotechnologie, Laval, Québec, Canada; Department of Biology, Faculty of Science and Medicine, University of Fribourg, Fribourg, 1700, Switzerland

## Abstract

Systemic immunity induced by beneficial bacteria helps plants to cope with subsequent pathogen attacks. However, how these bacteria are sensed by plants and how this sensing results in a systemic response is not well understood. Recently, the *Brassicaceae* LIPOOLIGOSACCHARIDE-SPECIFIC REDUCED ELICITATION (LORE) receptor kinase was identified as the direct sensor of bacterial medium-chain-length 3-hydroxy fatty acids (mc-3-OH-FAs) and 3-(3-hydroxyalkanoyloxy)alkanoic acids (HAAs), which activate systemic *Brassicaceae* immunity. Here, we show that the bacterium *Pantoea ananatis* BRT175, which produces HAAs, can colonize *Arabidopsis thaliana* roots independently of the production of these compounds. *P. ananatis* triggers induced systemic resistance (ISR) to the necrotrophic pathogen *Botrytis cinerea*, but not to the hemibiotrophic pathogen *Pseudomonas syringae* pv. *tomato*. Importantly, this ISR can be mimicked by HAAs or 3-hydroxy-decanoic acid (3-OH-C_10_). Both *P. ananatis*-triggered and 3-OH-C_10_-triggered ISR against *B. cinerea* are mediated by LORE sensing, involve salicylic acid, jasmonic acid and ethylene signaling pathways, and activate the expression of similar defense genes in infected leaves. Thus, 3-OH-C_10_ and HAAs are perceived by a pattern recognition receptor (PRR) in the root and activate a local immune response in roots as well as systemic immunity in leaves. Our study demonstrates that these lipidic microbe-associated molecular patterns (MAMPs), produced by beneficial microorganisms, are necessary and sufficient to trigger rhizobacteria-driven ISR. It further highlights the role of these MAMPs in differentially activating systemic resistance against biotrophic and necrotrophic pathogens.

## Introduction

Plants face numerous pathogenic threats in their natural environment. Throughout their evolutionary history, they have developed a defense system to limit infections and diseases (Finkel *et al*. 2017; Yu *et al*. 2019). Mutualistic bacteria in the rhizosphere can trigger systemic resistance to pathogens (Pieterse *et al*. 2014; Vlot *et al*. 2021). The term ‘induced systemic resistance (ISR)’ refers to the resistance expressed in plant tissues distant from those initially in contact with the pathogen (De Kesel *et al*. 2021). Most beneficial bacteria inducing ISR are plant growth-promoting rhizobacteria, often belonging to genera such as *Bacillus, Pseudomonas, Enterobacter, Klebsiella, Azospirillum, Paenibacillus*, and *Pantoea* spp. (Bhat *et al*. 2022; Zhu *et al*. 2022). In plants, activation of an immune response first relies on the perception of specific patterns, including Microbe-Associated Molecular Patterns (MAMPs), that are usually sensed by cell-surface pattern recognition receptors (PRRs), including receptor-like kinases (RLKs) or receptor-like kinase proteins (RLPs) (Boutrot and Zipfel, 2017; Ngou *et al*. 2022a; Zhang *et al*. 2023; Dodds *et al*. 2024). This MAMP-triggered immunity restricts pathogen proliferation, but also excessive colonization by beneficial microorganisms (Zamioudis and Pieterse, 2012; Wan *et al*. 2019; Ngou *et al*. 2022b). Although MAMPs from pathogenic as well as beneficial bacteria, like flagellin/flg22 or elf18, are recognized at the root level (Millet *et al*. 2010; Ranf *et al*. 2011; Wyrsch *et al*. 2015; Stringlis *et al*. 2018; Zhou *et al*. 2020), the interplay between MAMPs perception in roots and ISR remains poorly understood. Moreover, the involvement of specific PRRs in triggering ISR against pathogens with different lifestyles is not well documented. Following perception of microbes, ISR activation is regulated by hormonal signaling balance (Vlot *et al*. 2021). Roles of salicylic acid (SA), jasmonic acid (JA) and ethylene (ET) pathways, three of the key phytohormones involved in plant immunity, are still under debate, depending on the triggering microorganism, plant species and pathogens (Bürger and Chory, 2019).

Some beneficial strains of *Pantoea* are well-known for associating with plants (Coutinho and Venter, 2009) and for activating ISR (Walterson and Stavrinides, 2015; Duchateau *et al*. 2024). For instance, *Pantoea agglomerans* PTA-AF2 triggers ISR in grapevine against the fungus *Botrytis cinerea* (Trotel-Aziz *et al*. 2008; Verhagen *et al*. 2011), while *Pantoea ananatis* PS27 promotes growth and ISR against *Xanthomonas axonopodis* pv. *vesicatoria* in pepper (Kang *et al*. 2007). Strain BRT175 of *P. ananatis* is an epiphytic isolate from strawberries that produces effective antibiotics against the fire blight pathogen *Erwinia amylovora* (Smith *et al*. 2013). This strain synthesizes 3-(3-hydroxyalkanoyloxy)alkanoic acids (HAAs) as constitutive bricks of ananatoside glycolipids, a group of amphiphilic biosurfactants secreted by the bacterium (Smith *et al*. 2016; Gauthier *et al*. 2019). HAAs are synthesized by dimerization of (*R*)-3-hydroxyalkanoyl-CoA, derived from medium-chain 3-hydroxy fatty acids (mc-3-OH-FAs), through the RhlA enzyme (Abdel-Mawgoud *et al*. 2014). HAAs and mc-3-OH-FAs are perceived by the *Brassicaceae* S-domain receptor kinase LIPOOLIGOSACCHARIDE-SPECIFIC REDUCED ELICITATION (LORE) (Kutschera *et al*. 2019; Schellenberger *et al*. 2021). Sensing of these ligands by LORE triggers a canonical immune signature, leading to enhanced resistance against *Pseudomonas syringae* pv. *tomato* DC3000 (*Pst*) in *Arabidopsis thaliana* (thereafter Arabidopsis) leaves (Kutschera *et al*. 2019; Schellenberger *et al*. 2021). Interestingly, plants lacking LORE orthologs, like tomato, a *Solanaceae*, can gain mc-3-OH-FA immune sensing upon expression of *At*LORE (Eschrig *et al*. 2024). Here, we report that colonization of Arabidopsis roots by *P. ananatis* results in ISR against the necrotrophic pathogen *B. cinerea*. Although HAA synthesis is not a relevant trait for root colonization by *P. ananatis*, we demonstrate that HAAs and especially mc-3-OH-FAs mimic *Pantoea*-induced resistance. We show that *P. ananatis*-triggered and mc-3-OH-FA-triggered ISRs are mediated by the PRR LORE. Both ISRs require SA, JA and ET pathways and activate similar defense mechanisms in response to *B. cinerea*. Our data highlight the direct involvement of the PRR LORE and mc-3-OH-FA MAMP sensing in rhizobacteria-triggered ISR.

## Results

### *P. ananatis* colonizes Arabidopsis roots independently of HAA synthesis

We first assessed the ability of *P. ananatis* strain BRT175 to colonize Arabidopsis roots. Root inoculation with *P. ananatis* does not impede plant development nor provoke visible symptoms (Fig. 1A). Using a GFP-tagged *P. ananatis* BRT175 strain, we observed the colonization of the root system by the bacterium at seven days post-inoculation (dpi) (Fig. 1B). To investigate the role of HAAs in root colonization, we generated a GFP-tagged version of an *P. ananatis rhlA-*mutant, which is unable to synthesize HAAs (Smith *et al*. 2016). This tagged-mutant strain was also found in association with roots of Arabidopsis (Fig. 1B). By enumerating total bacteria at several time points (4 to 14 dpi), we found that both the WT and the mutant strains persist in the root system over the time course at stable titers between 10^5^ and 10^6^ CFU g FW^-1^ (Fig. 1C), showing that synthesis of HAAs does not impact colonization efficiency. Bacteria were also re-isolated from surface-sterilized roots, indicating their ability to colonize inner root tissues (Fig. 1C). No significant differences were observed between the WT and the *rhlA-*mutant of *P. ananatis* in this endophytic colonization. We did not retrieve bacteria from leaves of root-inoculated plants even after several days, indicating that *P. ananatis* colonization is restricted to the rhizosphere.

**Fig. 1:**
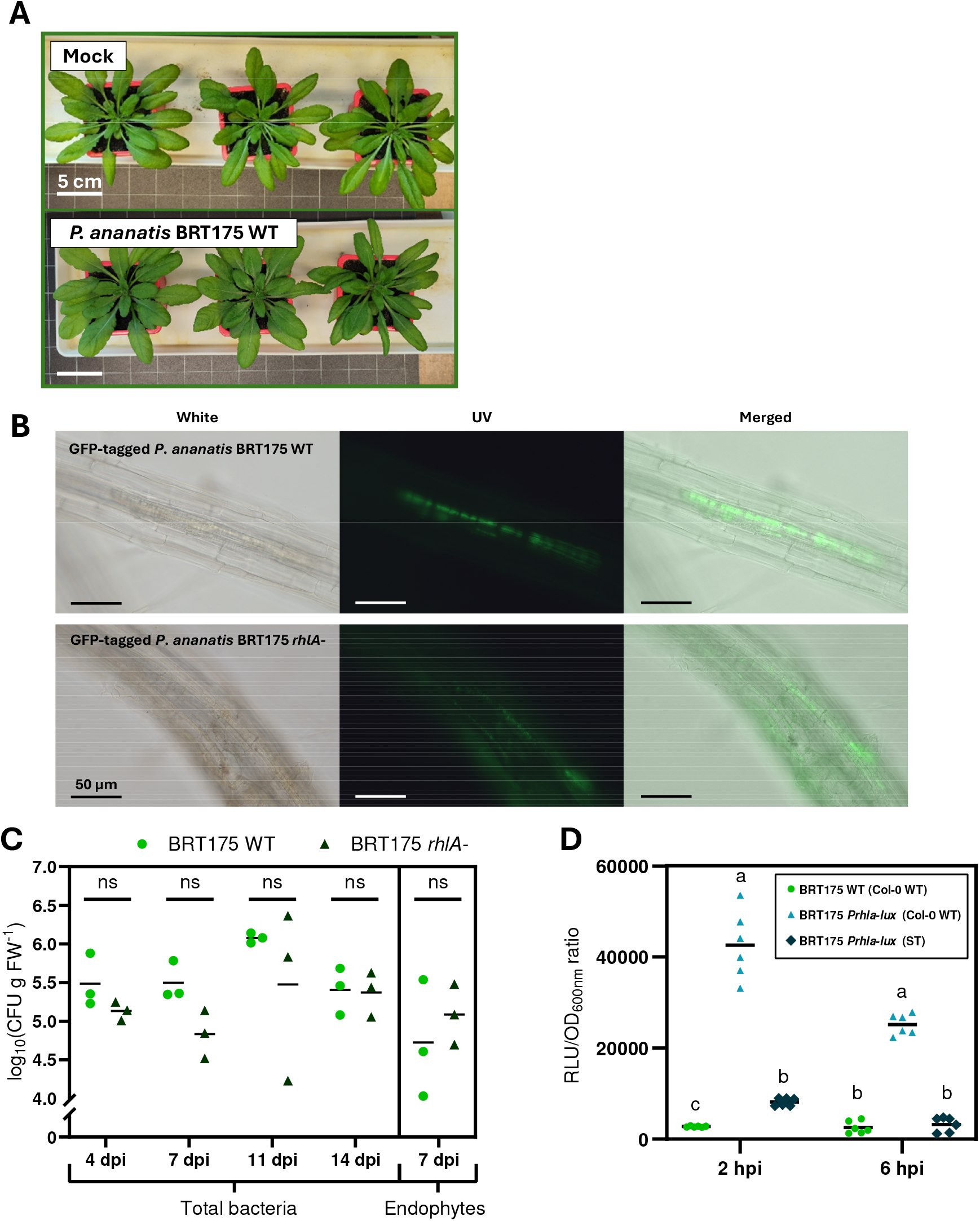
*P. ananatis* BRT175 colonizes Arabidopsis roots, and its association with plant tissues induces *rhlA* expression. **(A)** Pictures of mock-treated and root-bacterized *A. thaliana* Col-0 WT rosettes. Five-week-old plants were root-treated with 10 mM MgSO_4_ (Mock) or *P. ananatis* BRT175 WT (10^8^ CFU g of soil^-1^). Photos were taken two weeks after treatment. **(B)** Microscopic images of *A. thaliana* Col-0 WT roots colonized by GFP-tagged *P. ananatis* BRT175 WT or *rhlA-*mutant. Five-week-old plants were root-treated with a suspension of GFP-tagged *P. ananatis* BRT175 WT or *rhlA-*mutant (10^8^ CFU g of soil^-1^). Roots were unearthed from potting soil at 7 dpi and directly observed with a fluorescence microscope. **(C)** Enumeration of *P. ananatis* BRT175 WT and *rhlA-*mutant in unsterilized (total bacteria) and surface-sterilized roots (endophytes) of *A. thaliana* Col-0 WT. Five-week-old plants were root-treated with a suspension of *P. ananatis* BRT175 WT or *rhlA-*mutant (10^8^ CFU g of soil^-1^). Bacteria were re-isolated from roots at 4, 7, 11, and 14 dpi on LBA medium supplemented with rifampicin (50 mg L^-1^) for unsterilized roots and only at 7 dpi for surface-sterilized roots. Data are represented as individual values with means (black line) (n = 3). No significant differences were observed between both bacteria genotypes based on a non-parametric Mann-Whitney test (p ≤ 0.05). **(D)** Luminescence of *P. ananatis PrhlA-lux* mutant in contact with Arabidopsis plantlets grown in liquid MS medium. To evaluate the effect of living plant tissue on *rhlA* promoter activity, the mutant bacteria was also inoculated in MS medium containing sterile toothpicks (ST). *P. ananatis* BRT175 WT was used as a negative control. Data are mean ± SD (n = 6). Experiments were conducted twice with similar results.

Using a *P. ananatis* strain containing a transcriptional P*rhlA*-*lux* (promoter of *rhlA* fused with *luxCDABE*) reporter to measure the expression from the *rhlA* promoter, we found that expression was strongly up-regulated when bacteria were exposed to living plant roots compared to bacteria exposed to sterile toothpicks, used as a non-living control matrix, over a 6-h time course (Fig. 1D). These data suggest that exposure to living plant tissues significantly enhances HAAs biosynthesis in *P. ananatis*. Altogether, these observations indicate that while this strain can colonize the root system of Arabidopsis, HAA synthesis does not play an important role in this process, even though HAA production is induced by the presence of the plant.

### *P. ananatis* triggers an ISR against *B. cinerea* but not *Pst* in Arabidopsis

We then investigated the ability of *P. ananatis* to trigger ISR to the necrotrophic fungus *B. cinerea* and the hemibiotrophic bacterium *Pst* in soil-grown Arabidopsis. We inoculated roots with *P. ananatis* by soil drenching two weeks prior to leaf infection with *B. cinerea* or *Pst. B. cinerea-*induced necrosis symptoms were significantly reduced in leaves of root-bacterized plants as compared to leaves of mock-treated plants (Fig. 2A and 2B). In contrast, root treatment with *P. ananatis* did not reduce the titer of *Pst* nor associated disease symptoms in leaves (Fig. 2C and 2D).

**Fig. 2:**
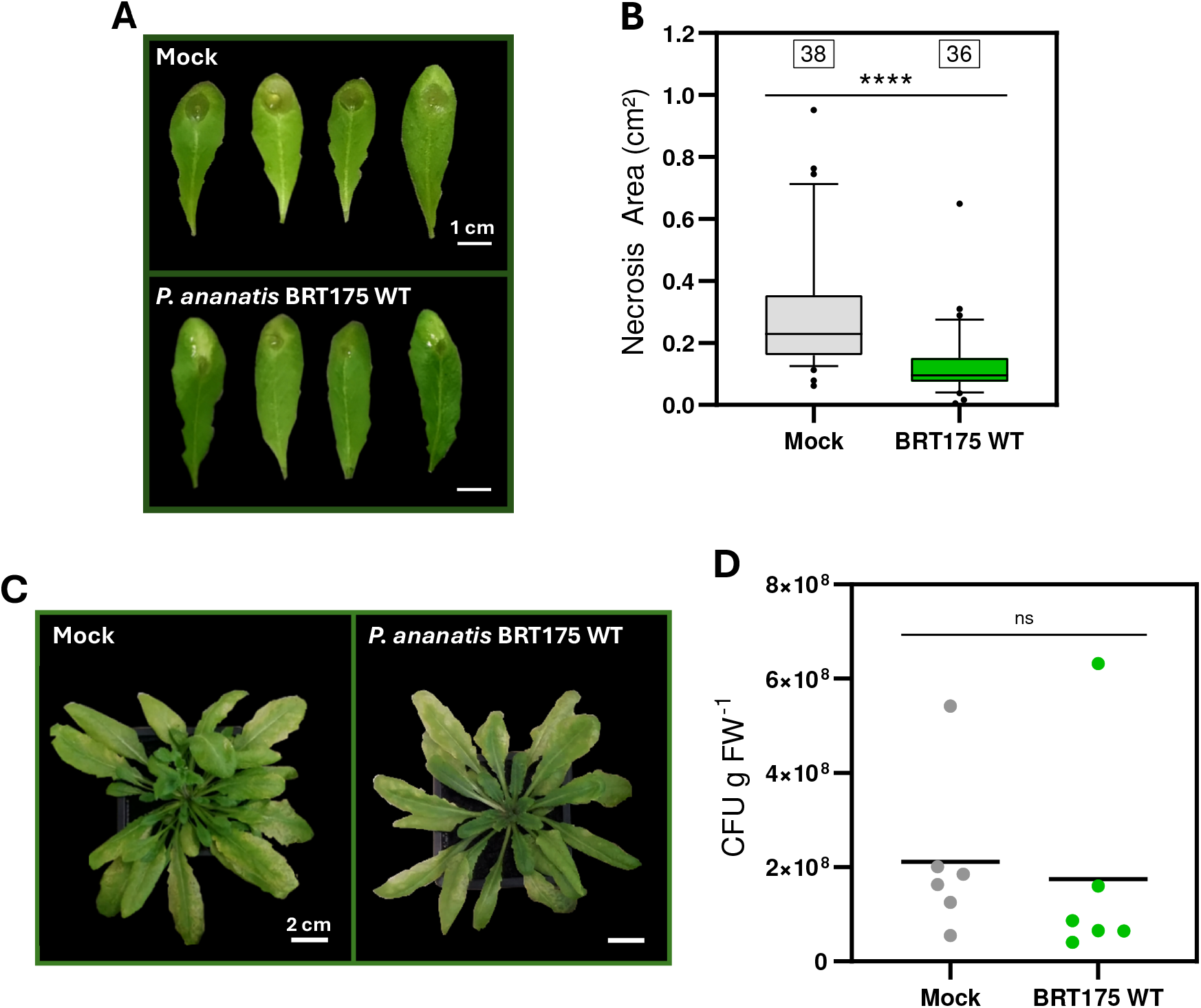
*P. ananatis* BRT175 induces a systemic resistance against *B. cinerea* but not against *Pst* in Arabidopsis. **(A)** Representative symptoms on Arabidopsis leaves caused by *B. cinerea* at 96 hpi. Five-week-old plants were root-treated with 10mM MgSO_4_ (Mock) or *P. ananatis* BRT175 WT (10^8^ CFU g of soil^-1^). Leaves were infected two weeks later with a 5 μL drop of *B. cinerea* spores (10^6^ spores·mL^-1^). **(B)** Necrotic symptoms provoked by *B. cinerea*. Lesions were measured at 96 hpi with ImageJ (n is indicated above each boxplot). **(C)** Representative symptoms on Arabidopsis leaves caused by *Pst* at 120 hpi. *A. thaliana* Col-0 WT were root-treated as previously described. Leaves were infected with *Pst* by spraying pathogens at 10^9^ CFU mL^-1^. **(D)** Enumeration of Colony-Forming Units (CFU) of *Pst*-infected leaves at 120 hpi. Data are represented individually (dots) with means (black line) (n = 6 different plants, for each, 6 foliar disks from 3 leaves are sampled and pooled). Asterisks indicate significant differences (p ≤ 0.05) based on a non-parametric Mann-Whitney test. Experiments were conducted three times with similar results.

### ISR triggered by *P. ananatis* against *B. cinerea* requires mc-3-OH-FA/HAA sensing by the PRR LORE

We then asked whether HAAs play a role in the ISR triggered by the beneficial bacterium. The *P. ananatis rhlA-*mutant strain induced less resistance against *B. cinerea* than the WT bacteria (Fig. 3A). Interestingly, HAAs alone induced a similar resistance phenotype as WT bacteria (Fig. 3A). Addition of HAAs to *P. ananatis rhlA-*during the inoculation process restored the ISR phenotype to the level observed with the WT strain (Fig. 3A). Since HAAs are sensed by the PRR LORE (Schellenberger *et al*. 2021), we tested the ability of *P. ananatis* WT to induce resistance in *lore*-5 mutant plants and in p*LORE:LORE* complemented *lore*-5 plants (Ranf *et al*. 2015). As shown in figure 3B, the *P. ananatis-*triggered ISR was lost in *lore*-5 plants but restored in p*LORE:LORE* complemented *lore*-5 plants. Additionally, we observed that both the *rhlA-*mutant strain and HAA were not able to trigger ISR in *lore*-5 plants (Supplemental Fig. 1).

**Fig. 3:**
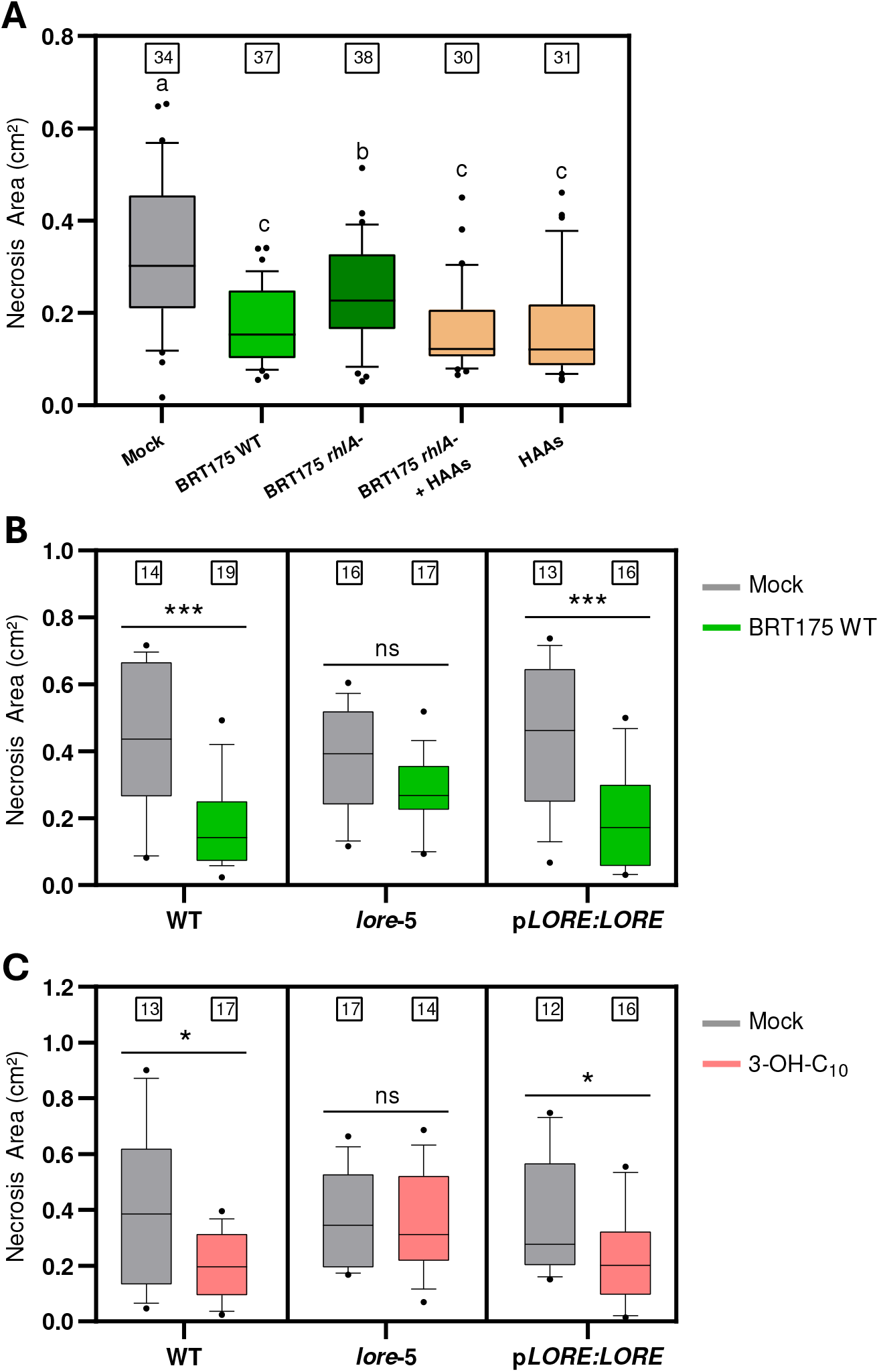
MAMP perception by the PRR LORE drives *P. ananatis* BRT175-triggered immunity in Arabidopsis against *B. cinerea*. **(A)** Necrotic symptoms provoked by *B. cinerea* in *A. thaliana* Col-0 WT. Plants were root-treated with 10mM MgSO_4_ (Mock), *P. ananatis* BRT175 WT or *P. ananatis* BRT175 *rhlA-*(both bacteria at 10^8^ CFU g of soil^-1^) or HAAs (directly supplied in bacterial suspension or in 10 mM MgSO_4_ at a final concentration of 500 μM). Leaves were infected two weeks later with a 5 μL drop of *B. cinerea* spores (10^6^ spores mL^-1^). Lesions were measured at 96 hpi with ImageJ. **(B)** Necrotic symptoms provoked by *B. cinerea* in *A. thaliana* Col-0 WT, *lore*-5 and p*LORE:LORE*, plants, root-treated and infected as previously described. Lesions were measured at 96 hpi with ImageJ. **(C)** Necrotic symptoms provoked by *B. cinerea* in *A. thaliana* Col-0 WT, *lore*-5 and p*LORE:LORE*, plants. Root were treated with MetOH 0.1 % (Mock) or with 3-OH-C_10_ (in 10 mM MgSO_4_ at a final concentration of 50 μM). Leaves were infected as previously described. Lesions were measured at 96 hpi with ImageJ. Different letters and asterisks indicate significant differences (p ≤ 0.05) based on a non-parametric Mann-Whitney test (n is indicated above each boxplot). All experiments were conducted three times with similar results.

We also investigated ISR activation against *B. cinerea* induced by 3-hydroxydecanoic acid [3-OH-C10:0] (thereafter 3-OH-C_10_), the strongest LORE agonist among the mc-3-OH-FAs group (Kutschera *et al*. 2019; Schellenberger *et al*. 2021). 3-OH-C_10_ triggers ISR against *B. cinerea* (Fig. 3C). This MAMP-mediated resistance was abolished in the *lore*-5 mutant but restored in p*LORE:LORE* complemented *lore*-5 plants (Fig. 4A). Like *P. ananatis*, 3-OH-C_10_ was unable to trigger ISR against *Pst* (Supplemental Fig. 2). Interestingly, 3-OH-C_10_ did not trigger resistance against *B. cinerea* when locally applied to leaves (Supplemental Fig. 3).

**Fig. 4:**
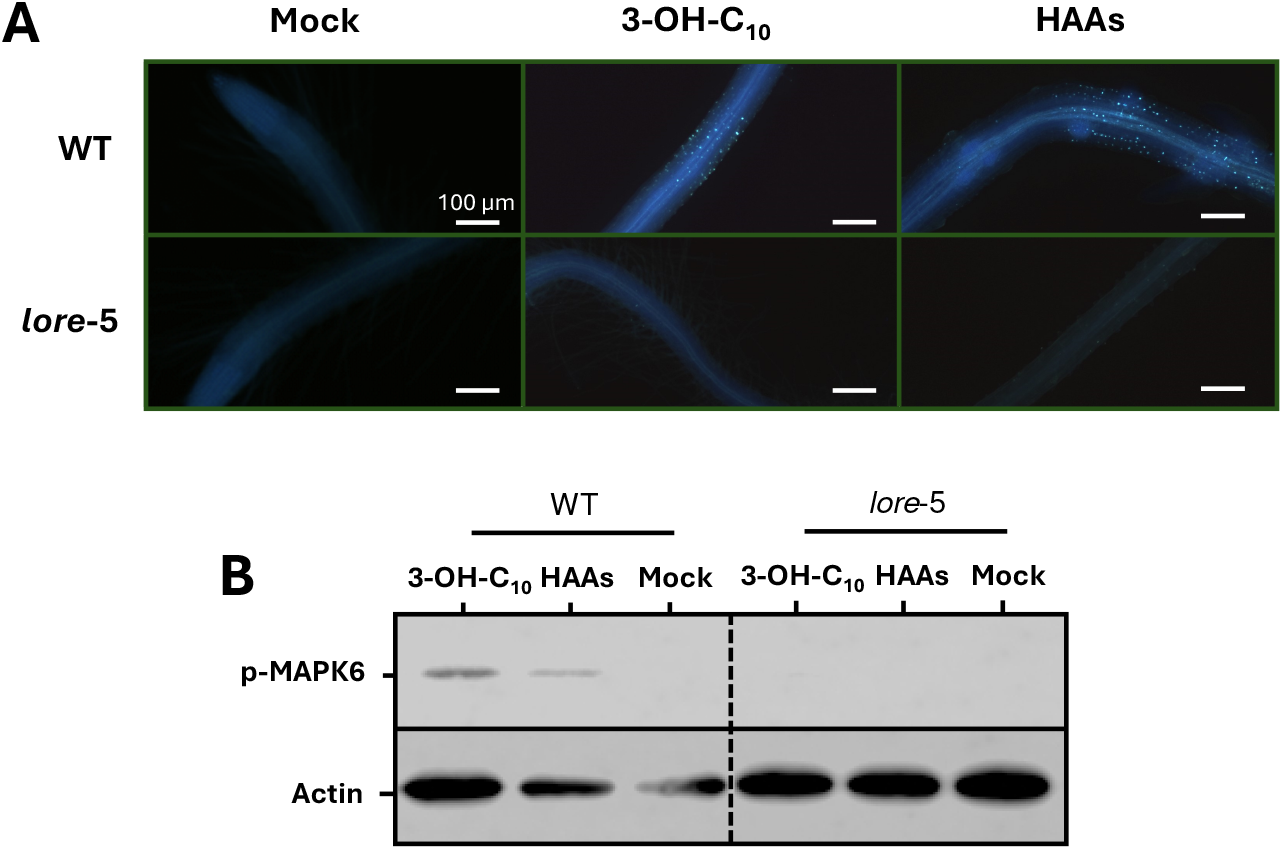
3-OH-C_10_ and HAAs activate root defense mechanisms in Arabidopsis through perception by the LORE receptor. **(A)** Callose deposits stained with aniline blue observed on treated roots of *A. thaliana* Col-0 WT or *lore*-5 plants. Roots were challenged with EtOH 0.1 % (Mock), 3-OH-C_10_ (10 μM) or HAAs (10 μM) and stained 24 hours later. **(B)** Phosphorylation of MAPK 6 in *A. thaliana* Col-0 WT and *lore*-5 plants. Roots were treated for 10 minutes with MetOH 0.1 % (Mock), 3-OH-C_10_ (1 μM) or HAAs (10 μM). Actin was used as a loading control. Full pictures are provided in Supplemental figure 4. Experiments were conducted twice with similar results.

We then investigated local defense responses in roots following challenge with 3-OH-C_10_ and HAAs as marker of local perception. As an event of late plant defense response, we assayed callose deposition (Millet *et al*. 2010) (Fig. 4A). In WT plants, we observed callose deposits in plant roots treated with 3-OH-C_10_ or HAAs, but not in mock-treated roots or *lore*-5 mutant plants. We also evaluated MAPK phosphorylation as a marker of early immune signaling (Kutschera *et al*. 2019) (Fig. 4B). Both 3-OH-C_10_ and HAAs triggered phosphorylation of MPK6 in WT, but not in *lore*-5 plants. These results demonstrate that HAAs and 3-OH-C_10_ are perceived by the PRR LORE at the root level, inducing a local immune response and activating ISR against *B. cinerea* in leaves. *P. ananatis* BRT175 (both WT and *rhlA-*mutants) also induce phosphorylation of MAPKs. Interestingly, we also observed MAPK activation by *P. ananatis* BRT175 in *lore*-5 mutant plants (Supplemental Fig. 4).

### *P. ananatis* and 3-OH-C_10_-triggered ISR require similar signaling pathways and trigger similar defense mechanisms

We aimed to decipher the contribution of hormone signal transduction pathways in *P. ananatis*-driven ISR. *P. ananatis*-triggered ISR against *B. cinerea* was compromised in *sid2*-1 and *npr1*-1 mutant plants (Fig. 5A), which are impaired in SA accumulation (Wildermuth *et al*. 2001) and signaling (Wu *et al*. 2012), respectively. ISR was also compromised in the *jar1*-1 mutant (Fig. 5A), deficient in JA-isoleucine synthesis (Staswick *et al*. 2002) and in the *ein2*-1 mutant (Fig. 5A), deficient in ET signaling transduction (Alonso *et al*. 1999). Altogether, these observations indicate that SA, JA and ET phytohormones contribute to *P. ananatis*-triggered ISR against the necrotrophic fungus. We then investigated whether these hormone signaling pathways are also involved in ISR activation after 3-OH-C_10_ sensing. Our results show that the ISR was indeed compromised in *sid2*-1, *npr1*-1, *jar1*-1 and *ein2*-1 mutant plants compared to WT upon pre-treatment of roots with 3-OH-C_10_ (Fig. 5B). To further compare *P. ananatis* and 3-OH-C_10_-triggered ISR, we quantified expression of defenses genes *AtPR1* and *AtPR4* in infected leaves of root-treated Arabidopsis. *AtPR1* is well known as a SA-dependent defense gene marker (Lebel *et al*. 1998; Vlot *et al*. 2009), whereas *AtPR4* expression is dependent on ET (Lawton *et al*. 1994; Broekaert *et al*. 2006). We observed that transcript levels of both *AtPR1* and *AtPR4* were significantly higher at 24 hpi in plants root-treated with *P. ananatis* (Fig. 5C) or 3-OH-C_10_ (Fig. 5D) in comparison with mock-treated plants. Both of these observations show that *P. ananatis*- and 3-OH-C_10_-triggered ISR involve similar hormone signaling pathways and potentiate similar defense mechanisms at the leaf level against *B. cinerea*.

**Fig. 5:**
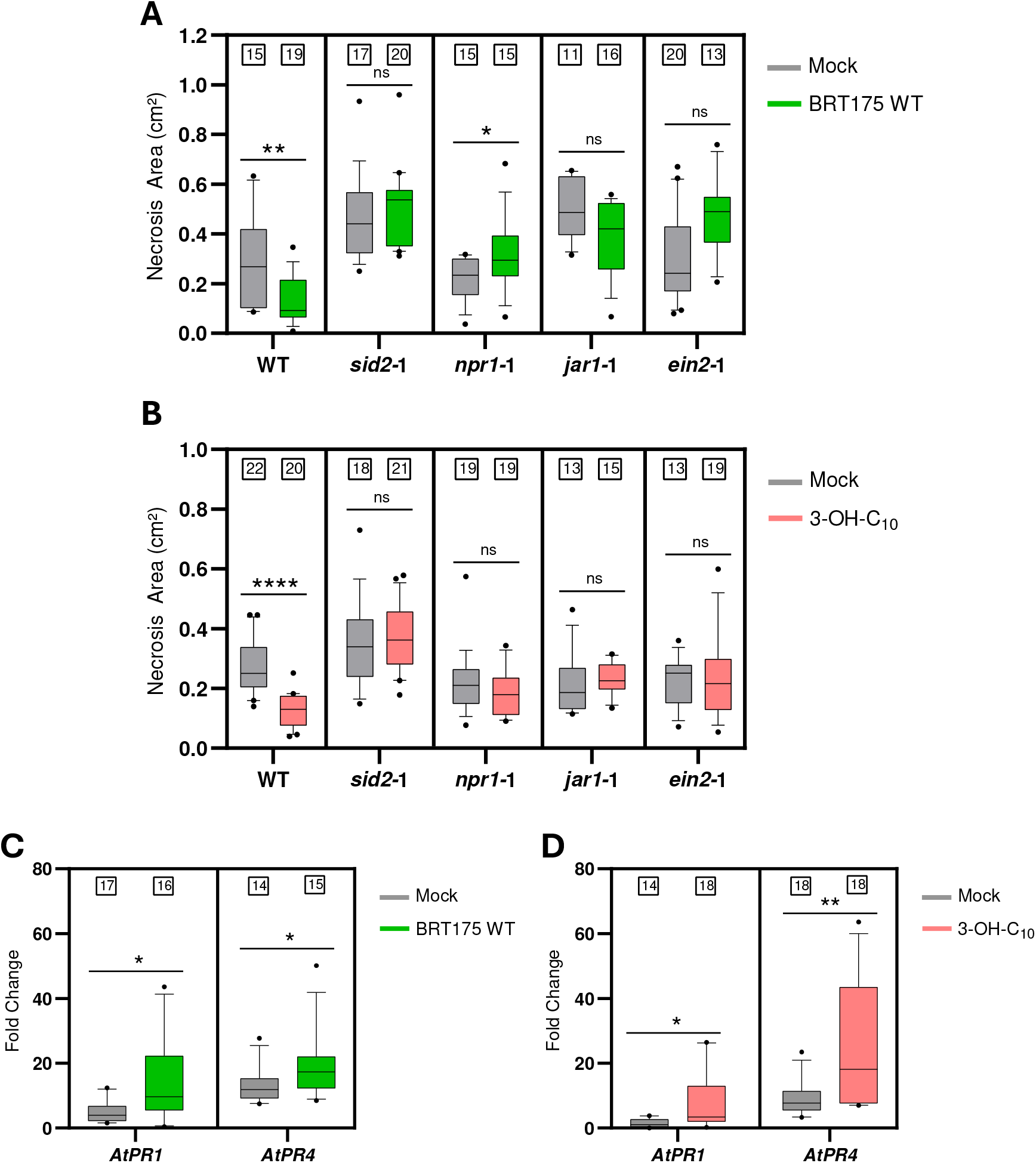
*P. ananatis* BRT175 and 3-OH-C_10_-triggered ISRs require identical signaling pathways and activates similar defense mechanisms against *B. cinerea*. Necrotic symptoms provoked by *B. cinerea* in *A. thaliana* Col-0 WT, *sid2-*1, *npr1-*1, *jar1-*1 and *ein2*-1 mutants, root-treated either with **(A)** 10mM MgSO_4_ (Mock) or *P. ananatis* BRT175 WT (10^8^ CFU g of soil^-1^), or with **(B)** MetOH 0.1 % (Mock) or 3-OH-C_10_ (in 10 mM MgSO_4_ at a final concentration of 50 μM). Leaves were infected two weeks later with a 5 μL drop of *B. cinerea* spores (10^6^ spores mL^-1^). Lesions were measured at 96 hpi with ImageJ. Different letters indicate significant differences (p ≤ 0.05) based on a non-parametric Mann-Whitney test (n is indicated above each boxplot). Experiments were conducted three times with similar results. Quantification of *AtPR1* and *AtPR4* expression in *A. thaliana* Col-0 WT leaves infected with *B. cinerea*. Plants were root-treated either with **(C)** 10mM MgSO4 (Mock) or *P. ananatis* BRT175 WT (10^8^ CFU g of soil-1), or with **(D)** MetOH 0.1 % (Mock) or 3-OH-C_10_ (in 10 mM MgSO4 at a final concentration of 50 μM). Leaves were infected with a spray of *B. cinerea* spores (10^5^ spores mL^-1^). Samples were collected at 0 and 24 hpi. For each timepoint and for each condition, 3 plants were samples (one sample consisting of 3 leaves pooled). Transcripts were quantified by qRT-PCR in triplicate for each samples. Results are expressed as fold change, calculated with the ΔΔCq method using *AtACT7* and *AtUBQ10* as housekeeping genes and mock-treated plants sampled at 0 hpi as the reference condition. Data are pooled from two independent biological replicates (6 different samples per condition in total, each quantified in triplicate). Asterisks indicate significant differences (p ≤ 0.05) based on a non-parametric Mann-Whitney test (n is indicated above each boxplot).

## Discussion

In this work, we first demonstrated that strain BRT175 of *P. ananatis* can colonize the root system of Arabidopsis as an endophyte. Many *Pantoea* bacteria can positively interact with plants (Walterson and Stavrinides *et al*. 2015; Duchateau *et al*. 2024). Some *P. ananatis* strains are characterized by their ability to colonize plant tissues (Kang *et al*. 2007; Gasser *et al*. 2012). We found that the lack of HAA biosynthesis does not affect the capacity of *P. ananatis* BRT175 to colonize the root system. Besides being released by some bacteria, HAAs are precursors for the synthesis of biosurfactants like rhamnolipids and ananatosides (Smith *et al*. 2016; Gauthier *et al*. 2019). Like their glycolipidic counterparts, HAAs play a direct role in swarming motility or adhesion to surfaces (Déziel *et al*. 2003; Smith *et al*. 2016). The role of microbial amphiphilic metabolites in plant colonization is dependent on the nature of the molecules. For instance, the cyclic lipopeptide massetolide A contributes to tomato colonization by *Pseudomonas fluorescens* (Tran *et al*. 2007). On the other hand, the lipopeptide poaeamide, synthesized by *Pseudomonas poae*, is a restrictive factor in this process (Zachow *et al*. 2015). Plant colonization by bacteria involves multiple factors. For example, the presence of pili/flagella on *P. ananatis* (Weller-Stuart *et al*. 2017) could participate in this process. Despite not playing a role in Arabidopsis root colonization, we provide evidence that HAA synthesis is induced when *P. ananatis* is exposed to Arabidopsis tissues, *via* activation of RhlA expression from the *rhlA* promoter. A positive effect of plant tissues on bacterial metabolite synthesis was also described for surfactin in *Bacillus* spp. (Nihorimbere *et al*. 2009; Debois *et al*. 2015; Hoff *et al*. 2021), syringopeptin (Wang *et al*. 2006) and amphisin (Koch *et al*. 2002) in *Pseudomonas* spp. Molecules such as small organic compounds, found in root exudates, could be involved in this process (Raaijmakers *et al*. 2010; Hoff *et al*. 2021).

Through its interaction with roots, *P. ananatis* activates a protection of Arabidopsis leaves against *B. cinerea*. While several strains of *P. ananatis* exhibit antagonistic features against plant pathogens (Torres *et al*. 2005; Gasser *et al*. 2012; Aman and Rai 2016), we found only one example of a systemic resistance inducer (Kang *et al*. 2007). As *P. ananatis* was not detected in leaf tissues, we postulate that this protection is solely driven by the induction of ISR upon sensing the bacteria or MAMPs they release in root tissues. We hence aimed to decipher the molecular actors involved in this ISR. As precursors of glycolipid biosynthesis, free HAAs are generally found in minimal concentrations in bacterial culture supernatants (Dubeau *et al*. 2009; Tiso *et al*. 2017; Germer *et al*. 2020). When synthesized by plant-associated bacteria, HAAs are thus most likely in contact with plant cells and membrane receptors. A recent study characterized the perception of HAAs by the PRR LORE, inducing Arabidopsis local immunity to pathogenic bacteria (Schellenberger *et al*. 2021). Using a *P. ananatis* mutant unable to synthesize HAAs, we demonstrated here that these lipids play a significant role in *P. ananatis*-mediated ISR against the necrotrophic fungus *B. cinerea*. In line with this observation, the external supply of HAAs to the *rhlA-*mutant strain restores the protection against *B. cinerea*. Concomitantly, we found that this ISR is fully dependent on bacterial perception involving the PRR LORE, as ISR was abolished in the *lore*-5 mutant plants, but restored in the complemented line. The partial protection observed in WT plants challenged with *P. ananatis rhlA-*could be explained by the perception of mc-3-OH-FAs by LORE (Kutschera *et al*. 2019). These molecules are also present in *rhlA-*bacteria, as a vital part of their primary lipid metabolism, and are potentially released by bacteria (Schellenberger *et al*. 2021). The role of MAMPs in bacterial-triggered ISR through PRR perception is not fully understood. Flagellin, for instance, is not a central actor of some bacterial-mediated ISR, although this peptide can trigger ISR when used as a purified compound (Meziane *et al*. 2005) and is also known for activating local root defenses (Millet *et al*. 2010; Zhou *et al*. 2020; Colaianni *et al*. 2021). Recently, the MAMP chitin was shown to drive a CERK1-dependent activation ISR to *Pst* in several plants, including Arabidopsis and lettuce (Makechemu *et al*. 2025). In line with our previous observations, we demonstrated that the MAMPs involved in *P. ananatis*-triggered ISR, HAAs and particularly mc-3-OH-FAs (here used in the form of 3-OH-C_10_), activate ISR against *B. cinerea* through sensing by the PRR LORE. We also showed that 3-OH-C_10_ and HAAs trigger MAPK phosphorylation and callose deposition in WT, but not in *lore*-5 Arabidopsis roots. Activation of LORE-dependent defense in roots by 3-OH-C_10_ has already been mentioned as 3-OH-C_10_ elicits defense responses in the elongation zone, but not in the differentiation zone of this organ (Zhou *et al*. 2020). *P. ananatis* BRT175 is supposed to present other MAMPs, like flg22 for instance (De Maayer and Cowan, 2016), which could also be sensed by plant cells. This could explain our observation that *P. ananatis* activate MAPK phosphorylation in roots of both WT and *lore*-5 mutant plants. Taken together, our results clearly demonstrate that LORE sensing of 3-OH-C_10_/HAAs is necessary and sufficient to trigger ISR to the necrotrophic fungus.

*P. ananatis* triggers ISR against the necrotrophic fungus *B. cinerea* but not against the hemibiotrophic bacterium *Pst*. Similarly, 3-OH-C_10_, albeit activating an ISR to the necrotrophic fungus, is unable to trigger ISR against *Pst*. The opposite effects on necrotrophic and hemibiotrophic pathogens may imply differential early or late signaling pathways post-perception. Interestingly, we found that *P. ananatis* and 3-OH-C_10_ mediated-ISR to *B. cinerea* involve SA, JA and ET pathways. Current models of signaling pathways in Arabidopsis immunity usually depict JA and ET as the most important phytohormones in ISR against necrotrophic pathogens while SA appears mostly involved in resistance against (hemi)biotrophic pathogens (Pieterse *et al*. 2014; Vlot *et al*. 2021). There is, however, some evidence that synergies of SA and JA pathways may occur in ISR, principally depending on the quantities and the timing of hormone production (Mur *et al*. 2006; Pieterse *et al*. 2012; Spoel and Dong, 2024). Moreover, our results are consistent with other studies showing that beneficial microorganisms, either bacteria (Nguyen *et al*. 2022) or fungi (Salas-Marina *et al*. 2011), can activate Arabidopsis ISR to necrotrophic pathogens *via* these two hormonal pathways. Regarding MAMPs, flg22 also triggers the accumulation of both phytohormones (Tsuda *et al*. 2008; Chang *et al*. 2017) and enhances the expression of genes involved in the SA and JA pathways (Denoux *et al*. 2008). *P. ananatis*- and 3-OH-C_10_-triggered ISR are both characterized by a potentiation of *AtPR1* and *AtPR4* expression in Arabidopsis leaves upon infection with *B. cinerea*. Activation of *AtPR1* is an outcome of an interplay between simultaneous low activations of the SA and the JA pathways (Mur *et al*. 2006; Loake and Grant, 2007). On the other hand, *AtPR4* expression depends on the JA and ET pathways (Lorenzo *et al*. 2003; Van Loon *et al*. 2006). Altogether, our data further emphasize the essential role of 3-OH-C_10_/HAAs in *P. ananatis*-triggered ISR, as we observed shared mechanisms between the MAMP-triggered and the bacteria-triggered defense response.

Based on previous studies (Kutschera *et al*. 2019; Schellenberger *et al*. 2021), we were also able to prove that 3-OH-C_10_ binding to LORE triggers different types of induced resistance depending on its site of perception. We found here that root perception of 3-OH-C_10_ activates ISR against the necrotrophic fungus *B. cinerea* but not against the hemibiotrophic bacteria *Pst*. Furthermore, leaf perception of this molecule does not trigger resistance against *B. cinerea*, while interestingly, previous studies conducted by Schellenberger *et al*. (2021) and Kutschera *et al*. (2019) showed an activation of local resistance against *Pst* in leaves through perception by the PRR LORE. Overexpression of this PRR in tomato leaves was also shown to enhance resistance against *Pst* (Eschrig *et al*. 2024). One explanation for this organ-dependent dichotomy may lie in the downstream signaling activated by LORE. To date, LORE is known to activate RIPK, involved in RBOHD activation (Wang *et al*. 2023), and the RLCKs PBL34, 35 and 36 (Luo *et al*. 2020). Further investigations into the direct and indirect targets of LORE could potentially reveal organ-related specificities in the signal transduction of mc-3-OH-FA sensing. Recently, LORE was also described as the key component governing perception of the pathogenic bacteria *Ralstonia solanacearum* and controlling xylem immunity in roots (Wang *et al*. 2023).

In conclusion, we demonstrate here that, when applied to Arabidopsis roots, *P. ananatis* BRT175 activates an ISR to the necrotroph *B. cinerea* but not to the hemibiotroph *Pst* in leaves. This ISR is solely driven by the perception of mc-3-OH-FAs/HAAs by the PRR LORE and involves SA, JA and ET signaling pathways (Fig. 6). As LORE sensing of 3-OH-mc-FAs/HAAs is activating a local resistance to *Pst* (Schellenberger *et al*. 2021) but not to *B. cinerea* (this study) in leaves, our work also highlights a differential effect on local/systemic resistance against necrotrophic/biotrophic pathogens following perception of a lipid MAMP by a PRR receptor.

**Fig. 6:**
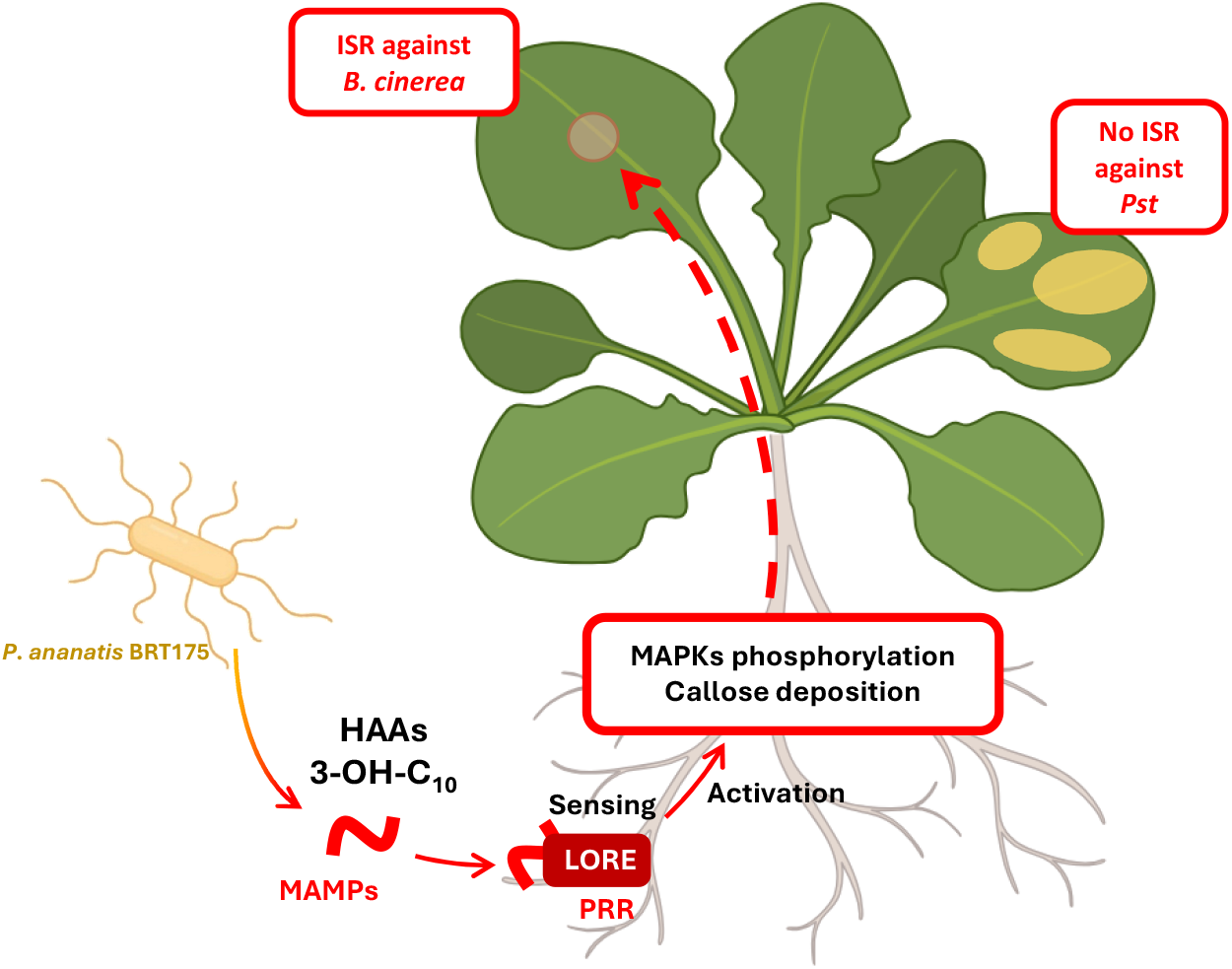
*P. ananatis* BRT175 triggers systemic immunity in Arabidopsis against *B. cinerea* through 3-OH-C_10_ and HAAs interaction with the PRR LORE. *P. ananatis* BRT175 synthetizes HAAs and 3-OH-C_10_. Root perception of these MAMPs by LORE activates root defense mechanisms. The defense signal reaches leaves, where it activates a defense response against *B. cinerea*. While triggering an ISR against a necrotrophic fungus, neither *P. ananatis* nor 3-OH-C_10_ mediated a systemic resistance against *Pst*.

## Material and Methods

### Plant material and growth conditions

*Arabidopsis thaliana* ecotype Columbia-0 (Col-0) was used in all experiments. In addition to the wild-type (WT) genotype, the *lore*-5, *sid2-*1, *jar1-*1, *npr1-*1 and *ein2*-1 mutants of the Col-0 ecotype (Schellenberger *et al*. 2021; Nguyen *et al*. 2022) and the *pLORE:LORE* complemented *lore*-5 line CLF12 (Ranf *et al*. 2015) were used in this study. All plants were grown in 50 g of potting soil (Terreau Motte, Sorexto) within a growth chamber (20°C, 12 h light/12 h dark, 150 μmol m^-2^ s^-1^, 60 % relative humidity). For the quantification of the *rhlA* promoter activity, plantlets were floating-cultivated in Murashige-Skoog (MS) medium. Seeds were first surface-sterilized as previously described (Millet *et al*. 2010), and disposed in 24-well plates, in MS medium with vitamins (Duchefa Biochemie) at pH 5.7 supplemented with sucrose (0.5 %) and MES monohydrate (0.05 %). Plates were sealed with Parafilm and incubated in a growth chamber under the same conditions. The medium was renewed after 8 days of incubation. For protection assays in hydroponics, plantlets were grown in Araponic systems. Seeds were placed in a seed-holder on a small plug of agar (0.6 %). The hydroponic fertilizer (Flora Series, Terra Aquatica) was added to the tray and changed every week. Araponic systems were placed in growth chamber (20°C, 12 h light/12 h dark, 150 μmol m^-2^ s^-1^, 60 % relative humidity).

### Microbial material and growth conditions

Both the wild-type (WT) *Pantoea ananatis* BRT175, isolated from a strawberry plant (Smith *et al*. 2013), and the isogenic *rhlA-*mutant (Smith *et al*. 2016), were used in this study. Bacteria were grown overnight in Lysogeny Broth (LB) medium at 28°C with shaking at 180 rpm. *P. ananatis* BRT175 is rifampicin-resistant, while *P. ananatis* BRT175 *rhlA-*is also kanamycin-resistant. When required, antibiotics were added to the liquid culture media at 50 mg L^-1^. For plant treatment, bacteria were collected by centrifugation (20 min, 3200 x *g*, 4°C) and suspended in sterile 10 mM MgSO_4_. Concentration of the bacterial suspension was adjusted to 10^9^ CFU mL^-1^.

*Botrytis cinerea Bc*630 was used as a necrotrophic pathogen (Nguyen *et al*. 2022). Conidia of *B. cinerea* were first germinated in Potato Dextrose Broth (PDB, 24 g L^-1^) for one week at 20°C under shaking (80 rpm). The mycelium from germinated conidia was then ground and spread on PDB Agar medium, followed by a two-week incubation at 20°C. Conidia were then harvested by scraping the mycelium with 4 mL of PDB. For plant infection, conidia were counted using a Malassez cell and the concentration was adjusted to 10^6^ conidia mL^-1^. Prior to infection, conidia were pre-germinated at 20°C under shaking (80 rpm) for two hours prior to the infection.

*Pseudomonas syringae* pv. *tomato* DC3000 (*Pst*) was used as a hemibiotrophic pathogen (Schellenberger *et al*. 2021; Nguyen *et al*. 2022). *Pst* was grown overnight in King’s B (KB) medium supplemented with rifampicin (50 mg L^-1^) at 28°C and 180 rpm. For infections, bacteria were collected by centrifugation (20 min, 3,200 x *g*, 4°C) and suspended in sterile 10 mM MgSO_4_ (10 mM). The concentration of the bacterial suspension was adjusted to 10^9^ CFU mL^-1^. Sterile Silwett-L77 was added to the bacterial suspension (0.02 %) as a wetting agent.

### Molecules

Purified 3-(3-hydroxyalkanoyloxy) alkanoic acids (HAAs) as C_10_-C_10_ congeners (Schellenberger *et al*. 2021) were used in this study. 3-Hydroxydecanoic acid (3-OH-C_10_) was purchased from Ark Pharm. All molecules were conserved at 100 mM in methanol (MetOH) or in ethanol (EtOH) at −20°C.

### GFP-tagging of *P. ananatis* BRT175 and its mutant

*P. ananatis* BRT175 (WT and *rhlA-*) were tagged with GFP to assess their ability to colonize root systems using epifluorescence microscopy. Competent cells were obtained from fresh cultures of both strains. Once the cultures reached an OD_600_ of 0.6, they were chilled on ice for 20 minutes. Pellets were collected by centrifugation (15 min, 16000 x *g*, 4°C) and washed twice with 20 mL of sterile water at 4°C. Pellets were then suspended in 4.5 mL of 10 % glycerol at 4°C and centrifuged again (10 min, 7500 x *g*, 4°C). Four milliliters of supernatant were discarded, and competent cells were suspended in the remaining 500 μL. Plasmid pIN301, carrying GFP and a chloramphenicol-resistance gene (Rabhi *et al*. 2018), was used to tag *P. ananatis* BRT175. 50 nanograms of plasmid were added to 100 μL of competent cells. The entire mixture was transferred to an electroporation cuvette (Gene Pulser/MicroPulser Electroporation Cuvettes, 0.2 cm gap, Bio-Rad). Electroporation was performed with an Eppendorf electroporator at 2.5 kV, 25 μF, and 200 Ω. Cells were immediately recovered after electroporation with 900 μL of LB medium supplemented with sterile dextrose (20 g L^-1^) and incubated for 2 h at 28°C with agitation at 180 rpm. Recovered cells were centrifuged (5 min, 3200 x *g*, 20°C) after incubation. 400 μl of supernatant were removed, and pellets were suspended in the remaining medium. 200 μl of suspended bacteria were spread on LB agar supplemented with chloramphenicol (50 mg L^-1^). After two days of incubation at 28°C, successful transformation was verified through direct observation of bacteria with an epifluorescence microscope (Olympus BH2) with UV light source (BH2-RFL-T3; Olympus).

### Root and leaf colonization assays

Five-week-old *A. thaliana* Col-0 WT plants were root-treated with different suspensions of *P. ananatis* BRT175 (WT or *rhlA-*mutant). Five milliliters of bacterial suspensions were inoculated directly into the soil, near the basis of the stem, to reach a final concentration of 10^8^ CFU g^-1^ of soil. Negative control plants were mock-treated with 5 mL of sterile 10 mM MgSO_4_ in a similar manner. Treated plants were placed in a growth chamber for a maximum of 14 days. Colonization was evaluated in roots and leaves at 4, 7, 11 and 14 days post-inoculation. At each time point and for each condition, six plants were collected. Whole plants were gently extracted from soil and roots were separated from the stem and leaves. The residual soil on the root system was removed with a gentle wash in water before sampling. To assess the endophytic capacity of both *P. ananatis* BRT175 strains (WT and mutant *rhlA-*), three out of the six roots were surface-sterilized, at each time point and for each condition. The sterilization process involved a general wash of roots with sterile MgSO_4_, followed by transfer to a diluted commercial bleach solution (0.5 % of active chlorine with 0.01 % Tween added) for 5 min. Sterilization ended with three 30 s washes of bleached roots with sterile MgSO_4_. All samples were ground in sterile mortars with 1 mL of sterile MgSO_4_ and diluted ten times. For each dilution, 10 μL were deposited twice on LB agar medium, supplemented with 50 mg L^-1^ rifampicin for selection of both phenotypes of *P. ananatis* BRT175. CFUs were counted after 2 days of incubation at 28°C. Results were expressed as a number of CFU per gram of root fresh weight. As negative controls, three root samples were collected from plants mock-treated with sterile MgSO_4_. Samples were ground using the same method and deposited on the same medium to assess the absence of rifampicin-resistant bacteria in Arabidopsis tissues. To control the efficiency of the sterilization process, 10 μL from the last wash with sterile 10 mM MgSO_4_ were deposited twice on the same medium.

### Microscopic observations of *A. thaliana* colonized roots

Following the same methodology described for colonization assays, GFP-tagged *P. ananatis* BRT175 were directly observed in the roots of Arabidopsis. Five-weeks old *A. thaliana* Col-0 WT plants were root-treated with a suspension of tagged-bacteria, and root systems were observed at 7 dpi. Roots were gently washed with water to remove remaining soil particles and were immediately observed using an epifluorescence microscope (Olympus BH2) with a UV light source (BH2-RFL-T3; Olympus). Roots were first observed under white light, and then under UV light. Both pictures were merged using the ImageJ software.

### Reporter gene construction

The activity from the *rhlA* promoter of *P. ananatis* BRT175 was evaluated by a fusion with the reporter genes *luxCDABE*. The *rhlA* promoter was amplified from genomic DNA of *P. ananatis* BRT175 (extracted using the Easy Pure Genomic DNA Kit, Trans) using the PaBRT175prhlA primers (sequences are listed in Supplemental Table I). PCR was conducted with the Q5® High Fidelity polymerase (New England Biolab) in a T100 Thermal Cycler (Bio-Rad). The PCR product was recovered from agarose using a FavorPrep Gel/PCR Purification Kit (Favorgen). The suicide vector pIJ514, containing a promoterless *luxCDABE* operon and a chloramphenicol-resistance gene within a Tn*7* transposon (Bessaiah *et al*. 2019), was linearized by digestion with EcoRI and SacI restriction enzymes (Thermo Scientific). The PCR product and linearized plasmid were assembled using the pEasy-Uni Seamless Cloning and Assembly Kit (Transgen) according to the manufacturer’s recommendations. The assembly product was introduced into competent *E. coli* SM-10 by heat shock. The construction was confirmed by PCR, using the same primers. After verification, the plasmid was transferred to competent *P. ananatis* BRT175 alongside the pSTNSK helper plasmid (Crépin *et al*. 2012) using electroporation. Chloramphenicol-resistant colonies of *P. ananatis* BRT175 were first selected at 30°C and then incubated at 37°C to lose the thermosensitive helper plasmid. Light emission of positive clones was verified using a Cytation multimode plate reader (BioTek).

### *rhlA* promoter activity assay

Evaluation of promoter activity was indirectly measured by monitoring the light emission of *P. ananatis* BRT175 *PrhlA-lux* cultivated alongside floating-cultivated *A. thaliana* plantlets. After 10 days of incubation, mutant bacteria were directly inoculated into MS medium at a final concentration of 10^8^ CFU mL^-1^. To compare the impact of living plant tissues on promoter activity, mutant bacteria were also inoculated in MS medium containing sterile toothpicks. Plant-bacteria co-culture medium was sampled (100 μL) at 2 and 6 h post-inoculation. Samples were placed in white 96-wells plate (OptiPlate, Perkin Elmer). Light emission and optical density at 600 nm were then measured directly using a CM SPARK (TECAN). Plantlets inoculated with *P. ananatis* BRT175 WT were used as negative control. For each condition, 6 wells (each containing approximately 15 plantlets) were inoculated. Data are expressed as the ratio between light emission and optical density.

### Bioprotection assays against *B. cinerea* and *Pst*

For the protection assays, plants were root-treated with bacterial suspensions as described for the colonization assays. However, these assays incorporated different genotypes of *A. thaliana* Col-0 (WT, *lore*-5, *pLORE:LORE, sid2*-1, *npr1*-1, *jar1*-1 and *ein2*-1). In addition to both *P. ananatis* BRT175 strains (WT or mutant *rhlA-*), plants were also root-treated with 3-OH-C_10_ or HAAs suspension, either alone or mixed with bacteria. 3-OH-C_10_ and HAAs were directly diluted (at 50 and 500 μM, respectively) in sterile 10 mM MgSO_4_ or in bacterial suspension for HAAs. After 2 weeks of incubation in a growth chamber, leaves of control and treated plants were infected as previously described (Nguyen *et al*. 2022). One day prior to infection, plants of similar conditions were placed in a plastic box containing 300 mL of tap water. Boxes were sealed with micropore adhesive tape and replaced in growth chamber for 24 hours. For *B. cinerea*, leaves were infected by placing a 5 μL drop of conidial suspension on the adaxial face of medium foliar-stage leaves. Infected plants were then replaced in sealed boxes. Infection was evaluated after 4 days of incubation in a growth chamber, by measuring the necrosis area provoked by *B. cinerea* using the ImageJ software. For each condition, approximately 15 to 30 leaves were infected on 6 different plants. For the infection with *Pst*, leaves were uniformly sprayed with bacterial suspension.

After 5 days of incubation in sealed boxes, 3 leaves were sampled for bacterial quantification. On each plant, 6 leaf discs (5 mm diameter) were excised from the 3 infected leaves and pooled together for grinding in sterile 10 mM MgSO_4_. Samples were then ten-fold diluted. Ten μL of the dilutions were spread on KB medium supplemented with rifampicin (50 mg L^-1^). CFUs were counted after 2 days of incubation at 28°C. Results are expressed as CFU per gram of fresh weight.

For protection assays conducted in hydroponic devices, 4-week-old plants were transferred to 10 mL vials containing hydroponic solution. Treatments were carried out by adding appropriate molecules (HAAs or 3-OH-C_10_) directly into the hydroponic solution, at respective final concentrations of 1 μM and 10 μM. 0.1 % EtOH was used as a negative control. After two days of incubation, leaves were infected with a drop of *B. cinerea* conidia suspension (10 μL at 10^5^ conidia mL^-1^). Necrosis areas were measured as described above. For the infection with *Pst*, leaves were uniformly sprayed with bacterial suspension (10^7^ CFU mL^-1^). Sterile Silwet L-77 was added to the bacterial suspension (0.02%) as a wetting agent. Three leaves were sampled and pooled per plant after 3 days of incubation in a growth chamber. CFUs were isolated and counted as described above.

For local protection assays, 5-week-old plants were leaf-treated by the spraying of a molecular suspension of 3-OH-C_10_ at 10 μM. EtOH 0.1 % was used as a negative control. After two days of incubation in a growth chamber, leaves were infected with a drop of *B. cinerea* conidia suspension (10 μL at 10^5^ conidia mL^-1^). Necrosis areas were measured as previously described.

### MAPK phosphorylation assays

MAPK phosphorylation state was measured in 2-week-old *A. thaliana* grown in hydroponic medium. The day prior to elicitation, twelve plantlets were incubated overnight in split Petri dishes to specifically treat the root systems. 3-OH-C_10_ (1 μM) or HAAs (10 μM) were added to root tips. MetOH 0.1 % was used as a negative control. Roots were then sampled and immediately placed in liquid nitrogen to preserve the phosphorylation state. The frozen roots were then ground. Protein extraction buffer (0.35 M Tris-HCl pH 6.8 containing 30 % (v/v) glycerol, 10 % (v/v) SDS, 0.6 M dithiothreitol, and 0.012 % (w/v) bromophenol blue) was then added (1 μL per 1 mg of root powder). Samples were then heated (10 min at 95°C) and centrifuged (10 min at 16000 x *g*). Supernatants were loaded into a polyacrylamide gel (SDS-PA gels 12 %). Proteins were separated by electrophoresis (15 min at 90 V, followed by 90 min at 150 V). Gel was then transferred to iBlot2 gel transfer system (Invitrogen) for protein migration on a PVDF membrane (7 min at 25 V). The membrane was then blocked using saturation buffer (3 % low-fat dry milk in Tris Buffered Saline (TBS)–Tween 20 (0.05 %)) for 30 min. Following 2 washes using washing buffer (0.5 % low-fat dry milk in TBS–Tween 20), the membrane was incubated overnight at 4°C with primary antibodies (Anti-phospho-p44/42 MAP kinase, Cell Signaling Technologies, diluted 1:2000 in washing buffer). The membrane was then washed 3 times and incubated for 1 h at room temperature with secondary antibodies (HRP-conjugated anti-rabbit, Bio-Rad, diluted 1:3000 in washing buffer). SuperSignal® West Femto (Thermo Fisher Scientific) was used as a revealing reagent. Luminescence was detected using Odyssey® Fc Dual-Mode Imaging System (LI-COR). Actin was revealed to normalize protein loading. The membrane was incubated for 30 min in 0.25 M NaOH solution, then blocked with saturation buffer. The membrane was incubated for 1 h at room temperature with primary antibodies (Plant monoclonal anti-actin, CusAb, 1:1000 in washing buffer), washed 3 times and incubated for 1 h at room temperature with secondary antibodies (Anti-mouse IgG HRP-conjugated, Cell Signaling, 1:3000 in washing buffer). The membrane was revealed as described above.

### Callose deposition assays

Callose deposits were observed in 2-week-old *A. thaliana* plantlets grown in hydroponics. 3-OH-C_10_ or HAAs were directly added into the culture medium at a final concentration of 10 μM. EtOH 0.1% was used as a negative control. After 24 h of treatment, root systems were sampled and placed in a discoloration solution (ethanol: acetic acid [3:1/v:v]) for 2 h. Roots were rehydrated in EtOH 70 % for 2 h, then in EtOH 50 % for 2 h, and finally in MilliQ water overnight. Subsequently, roots were incubated for 1 h at 37°C in NaOH 10 %, and in K_2_HPO_4_ 150 mM for 30 min, following several washes with MilliQ water. Roots were finally incubated in staining solution (K_2_HPO_4_ 150 mM and Aniline Blue 0.01 %) for 3 h in the dark. Stained roots were observed under an epifluorescent microscope (Olympus BH2) equipped with a UV light source (BH2-RFL-T3; Olympus).

### Quantification of defense gene expression

RNA extractions were performed on leaf samples of infected *A. thaliana* Col-0 WT. Soil-grown plants were root-treated with *P. ananatis* BRT175 WT or 3-OH-C_10_ and infected with *B. cinerea* as described above. Leaves were sampled at 0 and 24 hpi. For each time point and condition, 3 different plants were sampled (3 leaves were collected on each plant and pooled to make a unique sample per plant). Samples were freeze-dried in liquid nitrogen and ground to fine powder. Total RNA was extracted from 50 mg of ground tissue using QIAzol Lysis Reagent (Qiagen) according to the manufacturer’s instructions. RNA samples were reverse-transcribed using the Verso cDNA synthesis kit (Thermo Scientific) according to the manufacturer’s recommendations. To quantify gene expression, qPCR reactions were performed with the qPCRBIO SyGreen Blue Mix Lo-ROX (Eurobio Scientific, PCR Biosystem) on cDNA samples (1:10 diluted in RNAse-free water). Primer pairs were used at a final concentration of 0.3 or 0.1 μM, depending on the targeted gene. Sequences are listed in Supplemental Table I. PCR reactions were performed in white 384-well plate (Sorenson) with the CFX Opus 384 Real-Time PCR System (Bio-Rad) using the following procedure: first denaturation (15 min, 95°C), 40 cycles of denaturation (5 s, 95°C) and annealing/extension (30 s, 65°C), melting curve between 60°C and 95°C. Reactions were conducted in triplicate for each sample. Wells presenting a melting temperature difference of more than 0.5°C were excluded from the analysis (based on the melting curve of a PCR reaction performed on a pool of randomly selected samples). Expression of defense genes was normalized with the mean Cq of *AtACT7* and *AtUBQ10*, validated as housekeeping genes with the GeNorm algorithm. Expression was then calculated as fold change with the ΔΔCq method based on control samples (Mock-treated *A. thaliana* at 0 hpi).

### Statistical analysis

Statistical analyses were conducted using GraphPad Prism v10.6.1. Statistical differences of means were tested using Mann-Whitney tests. Differences were considered statistically significant at p ≤ 0.05.

## Acknowledgment

This work was supported by grants from the GLYCOBAC project, co-funded by the Région Grand Est and the ABIES Doctoral school. The authors also would like to gratefully acknowledge the MOBDOC program at the University of Reims Champagne-Ardenne and ABIES doctoral school for funding international mobility. Financial support from MESRI (Ministère de l’Enseignement Supérieur, de la Recherche et de l’Innovation) is also gratefully acknowledged. The authors also wish to thank C.M. Dozois (INRS-Centre Armand-Frappier Santé Biotechnologie, Laval, Québec, Canada) for providing plasmids pIJ514 and pSTNSK.

