## Supplemental Fig. 1 for "*Pantoea ananatis*-triggered systemic resistance requires root sensing through the LORE receptor kinase in Arabidopsis"

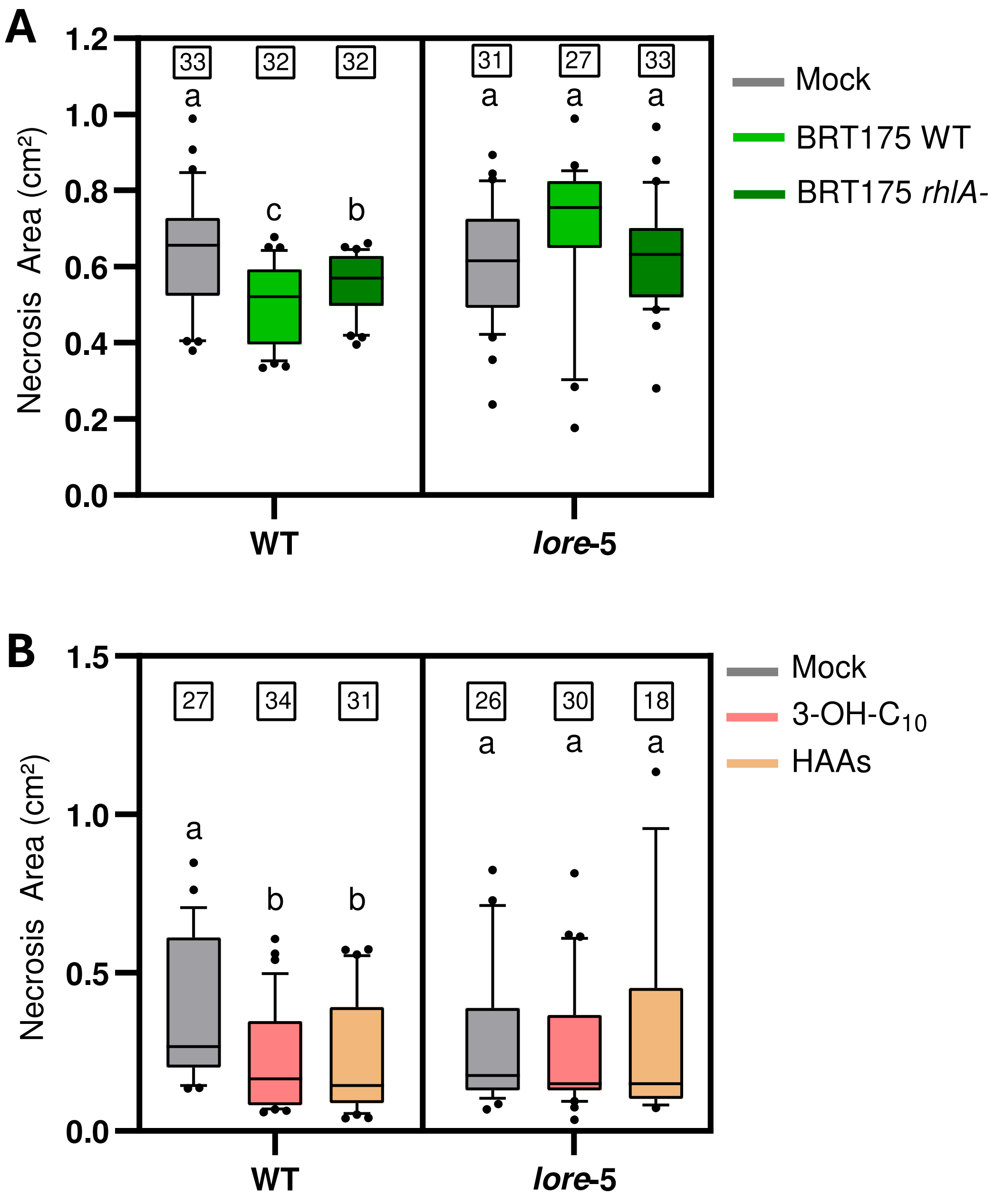


**Supplemental Data 1: ISR triggered by *P. ananatis* BRT175 *rhlA-* and HAA are abolished in *lore*-5 mutant Arabidopsis. (A)** Necrotic symptoms provoked by *B. cinerea* in *A. thaliana* Col-0 WT and *lore*-5 mutants, root-treated with 10mM $MgSO_4$ (Mock), *P. ananatis* BRT175 WT or *P. ananatis* BRT175 *rhlA-* (both bacteria at $10^8$ CFU g of soil$^{-1}$). Leaves were infected two weeks later with a 5 µL drop of *B. cinerea* spores ($10^6$ spores mL$^{-1}$). Lesions were measured at 96 hpi with ImageJ. **(B)** Necrotic symptoms provoked by *B. cinerea* in *A. thaliana* Col-0 WT and *lore*-5 plants. Arabidopsis were grown in hydroponics for four weeks. Roots were treated with EtOH 0.1% (Mock), 3-OH-$C_{10}$ (10 µM) or HAAs (1 µM) directly supplied in MS medium. Leaves were infected two days after treatment with a 10 µL drop of *B. cinerea* spores ($10^5$ spores mL$^{-1}$). Lesions were measured at 72 hpi with ImageJ. Different letters indicate significant differences ($p \leq 0.05$) based on a non-parametric Mann-Whitney test (n is indicated above each boxplot). Experiments were conducted twice with similar results.
