## Supplemental Fig. 2 for "*Pantoea ananatis*-triggered systemic resistance requires root sensing through the LORE receptor kinase in Arabidopsis"

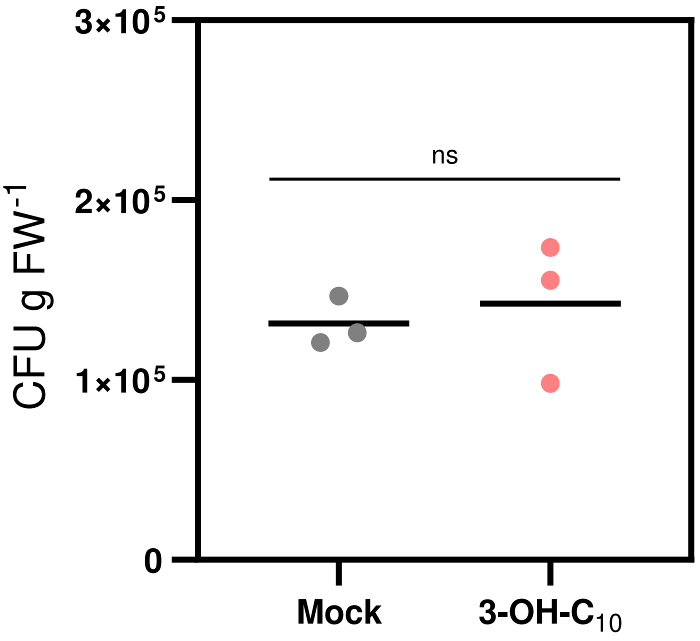


**Supplemental Data 2: 3-OH-$C_{10}$ does not trigger an ISR against *Pst* in Arabidopsis.** Enumeration of Colony-Forming Units (CFU) of *Pst* infected leaves at 72 hpi. *A. thaliana* Col-0 WT were grown in hydroponics for four weeks. Roots were treated by EtOH 0.1 % (Mock) or 3-OH-$C_{10}$ (10 µM), directly supplied in MS medium. Leaves were infected with *Pst* by spraying pathogens at $10^7$ CFU $mL^{-1}$. Data are represented individually (dots) with means (black line) (n = 3 different plants, for each, leaves were sampled and pooled). Asterisks indicate significant difference (p ≤ 0.05) based on a non-parametric Mann-Whitney test. Experiments were conducted twice with similar results.

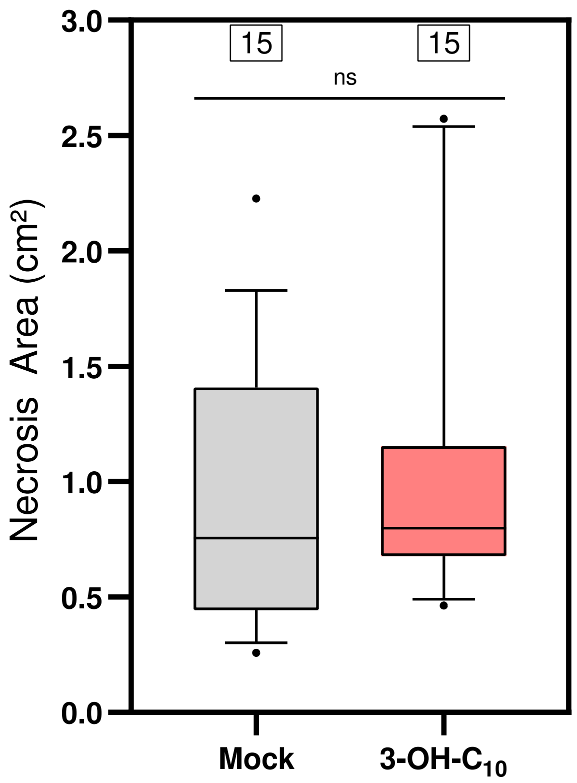


**Supplemental Data 3: 3-OH-$C_{10}$ does not trigger a local resistance against *B. cinerea* in Arabidopsis leaves.** Necrotic symptoms provoked by *B. cinerea* in *A. thaliana* Col-0 WT. Plants were leaf-treated with EtOH 0.1 % (Mock) or 3-OH-$C_{10}$ (10 µM). Leaves were infected two days later with a 10 µL drop of *B. cinerea* spores ($10^5$ spores $mL^{-1}$). Lesions were measured at 96 hpi with ImageJ (n is indicated above each boxplot). Asterisks indicate significant difference (p ≤ 0.05) based on a non-parametric Mann-Whitney test. Experiments were conducted twice with similar results.
