## Supplemental Fig. 4 for "*Pantoea ananatis*-triggered systemic resistance requires root sensing through the LORE receptor kinase in Arabidopsis"

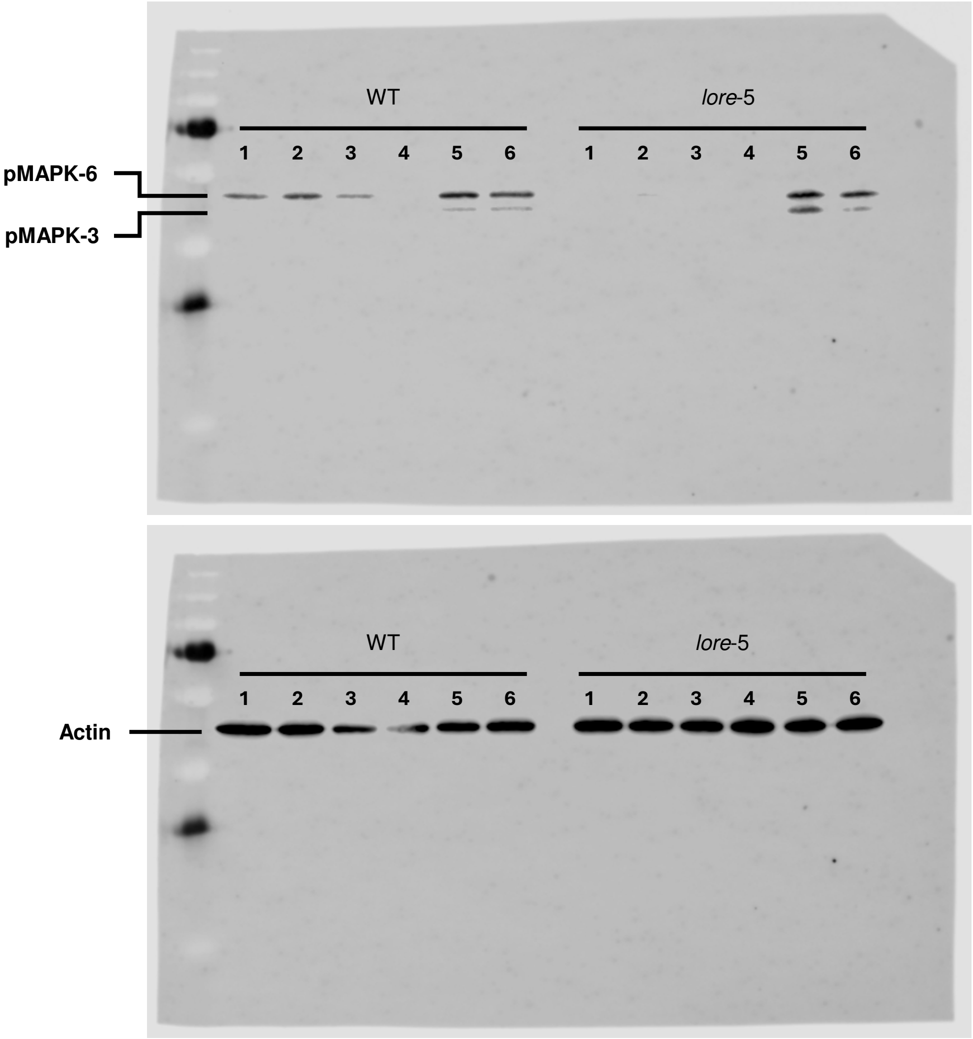


**Supplemental Data 4: Full-length blots of figure 4.** 1 and 2 : 3-OH-$C_{10}$ (1 µM), 3 : HAAs (10 µM), 4 : MetOH (0.1 %), 5 : *P. ananatis* BRT175 WT ($10^8$ CFU $mL^{-1}$), 6 : *P. ananatis* BRT175 *rhlA*- ($10^8$ CFU $mL^{-1}$).
