## Supplemental Table I for "*Pantoea ananatis*-triggered systemic resistance requires root sensing through the LORE receptor kinase in Arabidopsis"

**Supplemental table I: List of primers used in this study**

List of primers used for qRT-PCR assays

| Gene | Forward sequence (5'->3') | Reverse sequence (5'->3') | Concentration | Locus |
|---|---|---|---|---|
| *AtPR1* | AACTACGCTGCGAACACGTG | TCACTTTGGCACATCCGAGTC | 0.3 µM | AT2G14610 |
| *AtPR4* | AACAATGCGGTCGTCAAGGC | AAGCACTCACGGCTCTCAAATCCC | 0.3 µM | AT3G04720 |
| *AtACT7* | CCCAGGAATTGCTGACCGTA | TTTTCTCTCTGGCGGTGCA | 0.3 µM | AT5G09810 |
| *AtUBQ10* | GGTTTGTGTTTTGGGGCCTTG | CGAAGCGATGATAAAGAAGAAGTTCG | 0.1 µM | AT4G05320 |

List of primers used for promoter cloning

| Name | Forward sequence (5'->3') | Reverse sequence (5'->3') |
|---|---|---|
| PaBRT175prhlA | AATTCGATCATGCATGAGCTCTTATTATGCGCTGCAAAC | TGGGGCCCACCTCCTCGAGGTATCACTCTCCTTACCAAC |
